# DegQ and SwrA differentially modulate protease production and flagellar motility in *Bacillus subtilis*

**DOI:** 10.64898/2026.10.26.762088

**Authors:** Emily S. Brovey, Jacob A. Katz, Daniel B. Kearns

## Abstract

In *Bacillus subtilis,* the DegS-DegU two-component system modulates a range of adaptive responses, including extracellular protease production and the flagellar-surface motility called swarming. Two proteins, DegQ and SwrA, are thought to increase the amount of phosphorylated DegU and alter DegU DNA-binding abilities of DegU, respectively. Here we compare the relative contributions of DegQ and SwrA to DegU-related phenotypes in the *B. subtilis* ancestral strain that is functional for both proteins. We found that DegQ enhances the production of secreted proteases in stationary phase and inhibits swarming motility during exponential growth by a mechanism dependent on both DegS and DegU. Moreover, DegQ-mediated swarming inhibition was due, in part, to the attenuation of transcription levels within the long *fla/che* operon. SwrA by contrast, activates the promoter of the *fla/che* operon by a mechanism that depends on DegU but not DegS. Moreover, SwrA inhibits extracellular protease production by inhibiting the expression of DegQ. We conclude that DegQ and SwrA have opposing effects on DegU-dependent phenotypes and do so by different mechanisms. Thus, *B. subtilis* evolved third-party modulators to expand the conventional two-component system and achieve differential gene expression.

**IMPORTANCE:** The response regulator DegU regulates the expression of a wide array of adaptive responses including extracellular protease production and swarming motility in *B. subtilis*. Here we show that DegQ and SwrA modulate DegU activity to inversely regulate expression of the AprE protease and the *fla/che* operon for flagellar biosynthesis. This work highlights the roles of accessory factors in two-component system regulation, and clarifies the relationship between DegQ and SwrA, two proteins that were disabled during the domestication of laboratory strains.

## INTRODUCTION

*B. subtilis* secretes at least eight extracellular proteases, which while dispensable for growth in the laboratory, are likely necessary for nutrient acquisition in the environment (1, 2). Of the eight proteases, serine protease subtilisin E or AprE seems to be predominant and contributes to over 70% of the extracellular proteolytic activity in culture (3). Moreover, AprE has industrial applications with roles in leather and textile processing, food production, waste management, and pharmaceutical synthesis (4). Due to the biological consequences and extensive commercial applications, expression of *aprE*, the gene that encodes AprE, has been heavily studied and optimized (5–7). The primary regulator of *aprE* gene expression is thought to be the DegS-DegU two-component system (8).

DegU is a response regulator DNA-binding transcription factor that is phosphorylated by the soluble DegS kinase (9–12). Together, the DegS-DegU system is highly pleiotropic and regulates sporulation, biofilm formation (13, 14), genetic competence (15, 16), motility (17–19), poly-glutamate production (20, 21) and extracellular protease production (22–24) but discriminating direct from indirect effects is difficult due to the lack of a DegU DNA binding site consensus sequence (25, 26), and the input for the DegS kinase is unknown. Complicating matters further, DegU has been reported to activate and repress transcriptional activity in both its phosphorylated (DegU∼P) and non-phosphorylated states, and DegS-independent DegU activity and phosphorylation have been reported. Finally, DegU dependent gene expression is modulated by two small proteins, SwrA and DegQ, and both are defective in commonly studied laboratory strains of *B. subtilis* (13, 20, 27, 28).

The *swrA* gene that encodes SwrA contains a loss of function frameshift mutation in laboratory strains, and the first indication of SwrA function was the isolation of a mutant allele called *ifm*, that restored the reading frame to functionality and enhanced motility (27–29). Subsequent research indicated that SwrA increased the frequency of motile cells in the population (30) and increased the number of flagella per cell on motile individuals (30–32). ChIP-seq analysis indicated that SwrA enriched DNA in a DegU-dependent manner and did so at a subset of DegU regulated promoters (33). Moreover, SwrA induced DegU oligomerization at an activation sequence upstream of the *fla/che* operon dedicated to flagellar biosynthesis, allowing cells to upregulate transcription and reach the hyperflagellated state required for swarming (19, 33, 34). Despite SwrA enrichment at multiple DegU regulated promoters, what governs specificity is unknown, and SwrA does not currently have a clear role outside the regulation of motility.

The *degQ* gene encoding DegQ contains a mutation in the promoter that all but abolishes expression in laboratory strains, and *degQ* was originally discovered in a screen in which a reversion of this mutation enhanced extracellular protease production (20, 22, 35). Moreover, DegQ was shown to activate the promoter of *aprE* (P*_aprE_*) and does so in a DegU-dependent manner (6, 13, 36). DegQ is thought to modulate DegU phosphorylation, and either promotes the phospho-transfer from DegS, inhibits DegS phosphatase activity, or stabilizes the phosphorylated form of DegU∼P (13, 37). The precise mechanism by which DegQ stimulates DegU-dependent expression is unknown, and its role in regulating DegU-related phenotypes outside of extracellular protease production is poorly understood.

Here we explore the effect of DegQ and SwrA and their relative contribution to DegU-mediated gene regulation in the ancestral 3610 strain of *B. subtilis*. Whereas cells with high levels of DegQ increase production of extracellular protease activity but inhibit swarming motility. Conversely, SwrA is required for swarming but inhibits extracellular proteases. The phenotypic consequences of both proteins are reflected in their opposing effects on DegU-dependent expression of the P*_aprE_* and P*_fla/che_*. In the course of the study, we find that extracellular proteases are not expressed until the transition to stationary phase and that while the DegQ effect requires the DegS kinase, the SwrA effect does not. Moreover, we show that SwrA appears to act as an inhibitor of *aprE* gene expression by inhibiting the expression of DegQ. Thus, the two small protein modulators appear to act in opposition at the transition between exponential growth and stationary phase. How such different regulatory outcomes are achieved through the same DNA binding transcription factor remains unknown.

## RESULTS

### The DegS-DegU two-component system activates expression of *aprE*

To determine whether DegU was required for extracellular protease activity in the ancestral strain of *B. subtilis*, supernatants were collected over a time course and protease activity was measured. At low OD_600_ during exponential growth (**Fig 1A**), little protease activity was detected but increased at high optical density during the transition to stationary phase (**Fig 1B**). Moreover, disruption of DegU had no effect on growth (**Fig 1A**) but reduced extracellular protease activity relative to wild type at all optical densities tested (**Fig 1C**). We conclude that extracellular protease activity is a stationary phase phenomenon under the conditions tested and is enhanced by DegU in the ancestral strain of *B. subtilis*.

**Figure Legend 1.**
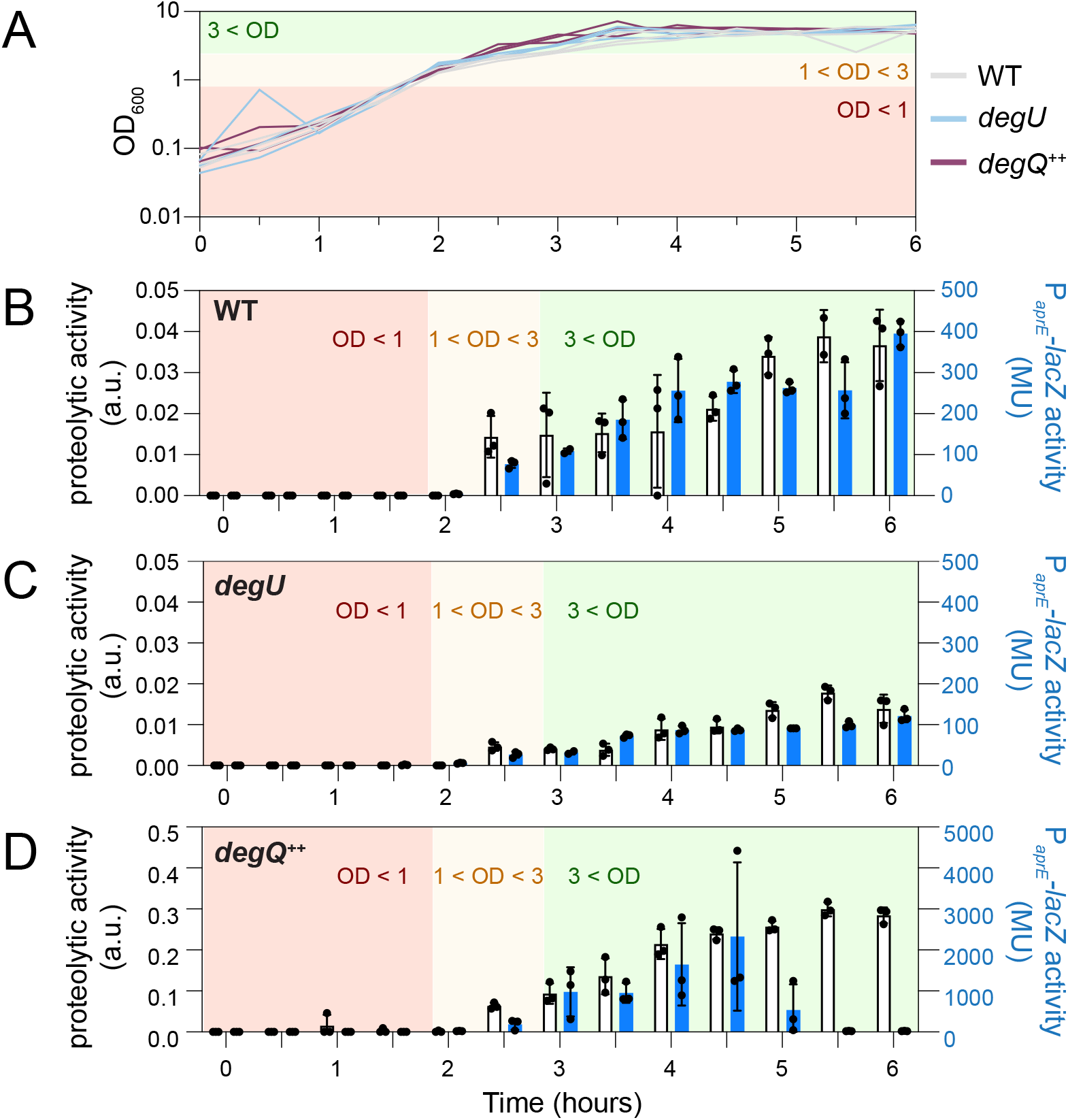
Extracellular protease production is activated by DegQ in stationary phase. A) Spectrophotometric growth curve of wild type and various strains measured at OD_600_ and grown in LB at 37 °C. Horizontal colored bars indicate regions of cell density where orange is less than 1 OD_600_, yellow is between 1 and 3 OD_600_, and green is above 3 OD_600_. The color coding is maintained throughout the remainder of the figure. The following strains were used to generate this panel: wild type (gray, DB3692), *degU* (blue, DB1815), and cells overexpressing *degQ*^++^ (pink, DB3695). B-D) Double Y bar graphs where white bars measure extracellular protease activity and blue bars measure the expression of a P*_aprE_*-*lacZ* transcriptional reporter. Each datapoint is the average of three replicates where the bar is the average, the whiskers represent the standard deviation, and the individual data points are shown as dots. The following strains were used to generate these panels and are the same as panel A: B) wild type (DB3692), C) *degU* (DB1815), and D) *degQ*^++^ (DB3695). To generate this figure, cells were grown to an OD600 of approximately 2, samples were collected, extracellular protease activity and β-galactosidase expression were measured in parallel

The alkaline protease AprE is encoded by the *aprE* gene, the expression of which is considered to be a primary indicator for proteolysis (3, 5), and disruption of AprE reduced the total amount of extracellular proteolytic activity (**Fig 2**). To measure *aprE* expression, a reporter construct in which the *lacZ* gene encoding β-galactosidase was fused to the P*_aprE_*promoter, inserted at an ectopic site, and measured at various optical densities. Consistent with the protease activity profile, expression of P*_aprE_* was low at low OD and increased during the transition to stationary phase (**Fig 1B**). Also consistent with previous observations, deletion of *degU* (**Fig S1A**) reduced expression from P*_aprE_* at all densities tested (**Fig 1C**). Based on these results, we chose an optical density of 2 for our standard conditions when measuring P*_aprE_* expression, as this was a condition with substantial activity that was reduced by mutation of *degU.* Moreover, the *degU* mutant phenotype could be complemented when *degU* was from its native promoters (10, 38), and inserted at an ectopic site in the chromosome (**Fig S1A, B**). Finally, deletion of *degU* resulted in a significant decrease in extracellular protease activity under the standard conditions (**Fig 2**).

**Figure Legend 2.**
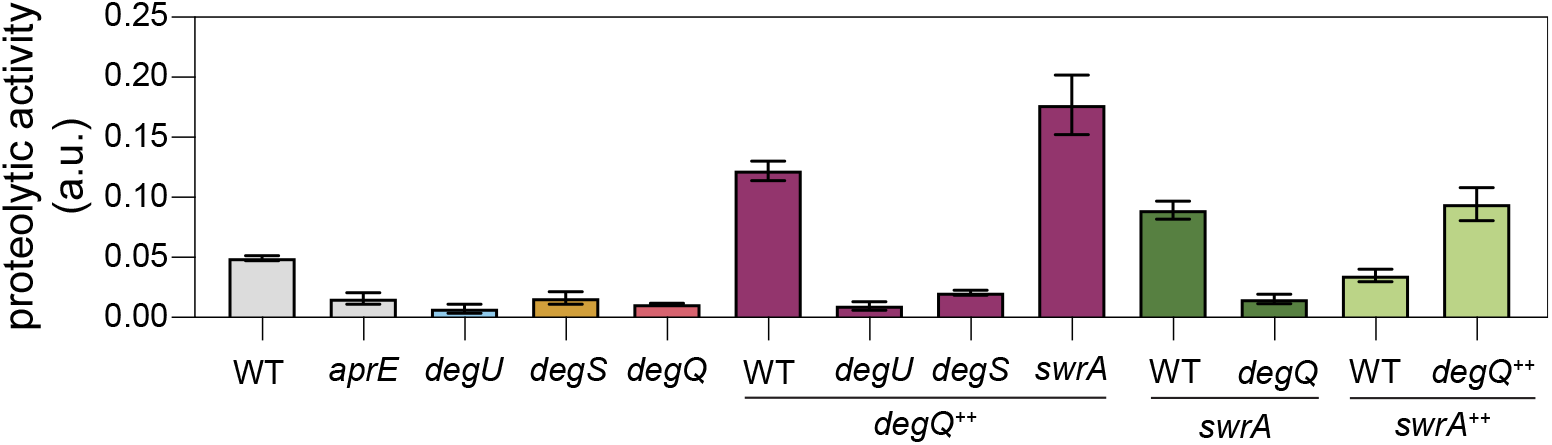
Extracellular protease activity is enhanced by DegQ and inhibited by SwrA. Extracellular protease activity as measured by the degradation of fluorescent azocasein in supernatants of cells grown to OD_600_ of ∼2 in LB (+1mM IPTG when appropriate) from strains of the indicated genetic backgrounds. Each bar is the average activity taken from at least three independent measurements, error bars represent the standard deviation. The dashed line indicates the amount of activity taken from wild type. The following strains were used to generate this panel: wild type (DK1042) *aprE* (DS5648), *degU* (DS3649), *degS* (DK4236), *degQ* (DB3074), *degQ^++^* (DK6249), *degU degQ^+^*^+^ (DB3745), *degS degQ^+^*^+^ (DB2863), *swrA degQ^+^*^+^ (DB3185), *swrA* (DS2415), *swrA*^++^ (DB3835), *swrA^++^ degQ^+^*^+^ (DB3729).

The DegU response regulator is phosphorylated, at least in part, by the cognate kinase DegS (11, 12, 39). To test whether the DegS kinase was required for the expression of *aprE,* the reporter construct was inserted into a strain deleted for the *degS* gene in a manner that preserved previously reported internal promoters that drive *degU* gene expression (38) (**Fig S1A**). The *degS* deletion significantly decreased expression of the *aprE* promoter and decreased extracellular protease activity (**Fig 2**, **Fig 3A**). Moreover, the mutation could be partially complemented by introduction of the *degS* gene expressed from its native promoter and inserted at an ectopic site in the chromosome (**Fig S1A, B**). We conclude that expression of *aprE* is dependent on both DegS and DegU and we infer that the phosphorylation of DegU is required for *aprE* expression.

**Figure Legend 3.**
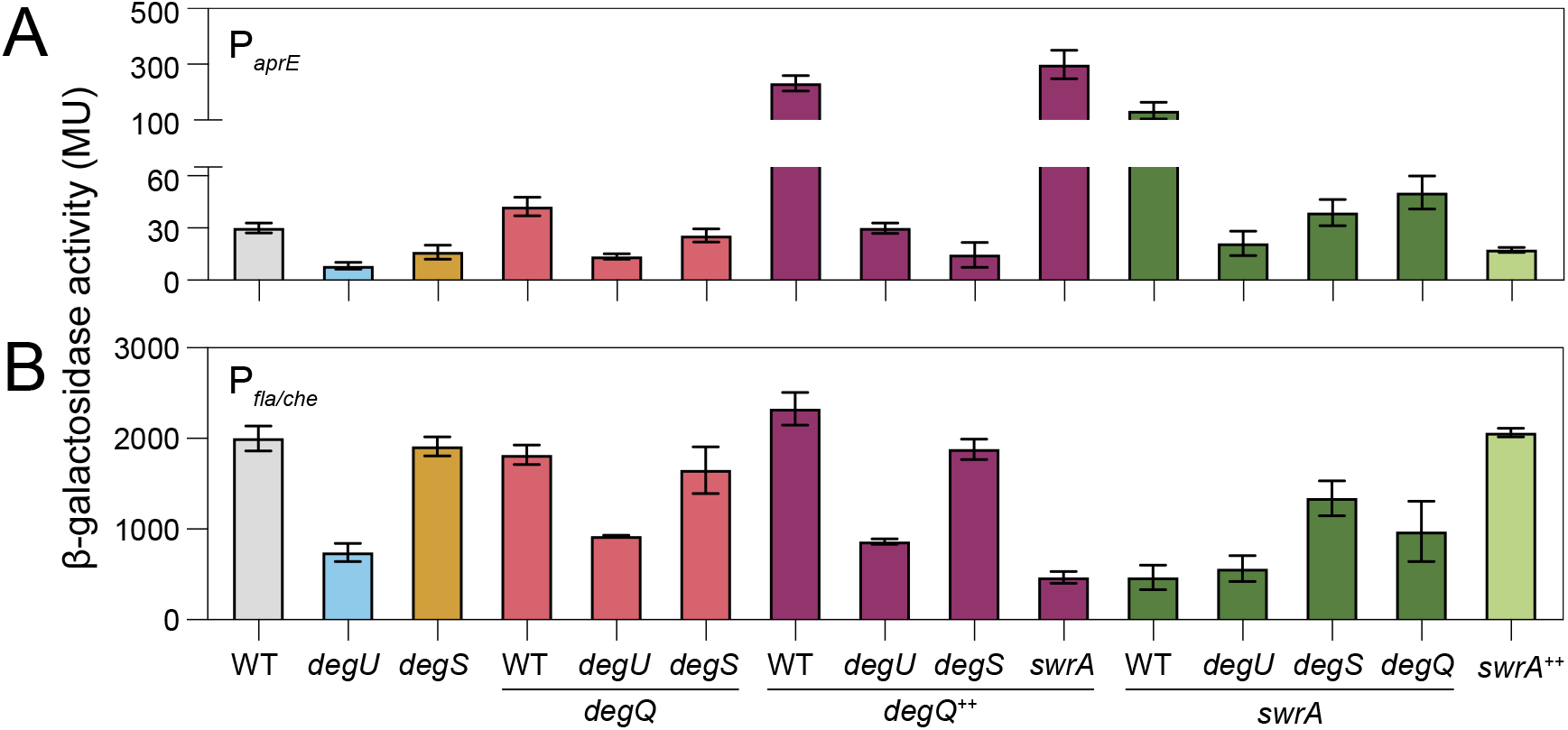
DegQ activates P*_aprE_* and SwrA inhibits DegQ expression. β-galactosidase activities measured in Miller units (MU) for each of the indicated transcriptional reporter fusions and mutants grown to an OD of 2 in LB media. A) Expression levels from the P*_aprE_* reporter. The following strains were used to generate this panel: wild type (DB3692), *degU* (DB1815), *degS* (DB3741), *degQ* (DB3751) *degU degQ* (DB3767), *degS degQ* (DB2877), *degQ^++^* (DB3695), *degU degQ^++^* (DB3769), *degS degQ^++^* (DB3693), *swrA degQ*^++^ (DB3748), *swrA* (DB3698), *degU swrA* (3750), *degS swrA* (DB3765), *degQ swrA* (DB3746), *swrA*^++^ (DB3865). B) Expression levels from the P*_fla/che_* reporter. The following strains were used to generate this panel: wild type (DK5183), *degU* (DB3697), *degS* (DB3742), *degQ* (DB3752) *degU degQ* (DB3768), *degS degQ* (DB3131), *degQ^++^*(DB3696), *degU degQ^++^* (DB3770), *degS degQ^++^* (DB3764), *swrA degQ*^++^ (DB3749), *swrA* (DK6345), *degU swrA* (DB3753), *degS swrA* (DB3766), *degQ swrA* (DB3747), *swrA*^++^ (DB3701). Each bar is the average of three biological replicates, and the error bars represent the standard deviation.

To determine whether the DegS-DegU two-component system activates transcription of the other extracellular proteases, RNA was purified from the wild type and cells deleted for either *degS* or *degU*, grown to an optical density of 2, sequenced, and compared in transcriptomic analysis. Consistent with β-galactosidase reporter assays, deletion of either *degS* or *degU* resulted in a decrease in the amount of transcript from the *aprE* gene and also reduced expression from gene *bpr* encoding the Bpr protease (**Fig 4A**) (26, 34). Transcripts from the 6 remaining genes that encode extracellular proteases, *epr*, *mpr*, *nprB*, *nprE*, *vpr*, and *wprA* were unchanged from wild type (**Fig 4A**) (25, 26). We conclude that DegS and DegU increase transcript abundance of *aprE* and *bpr*. While it does not appear that DegS and DegU regulate the remaining 6 extracellular protease encoding genes, we note that additional regulation may apply at those loci and potentially obscure the effect under our standard conditions.

**Figure Legend 4.**
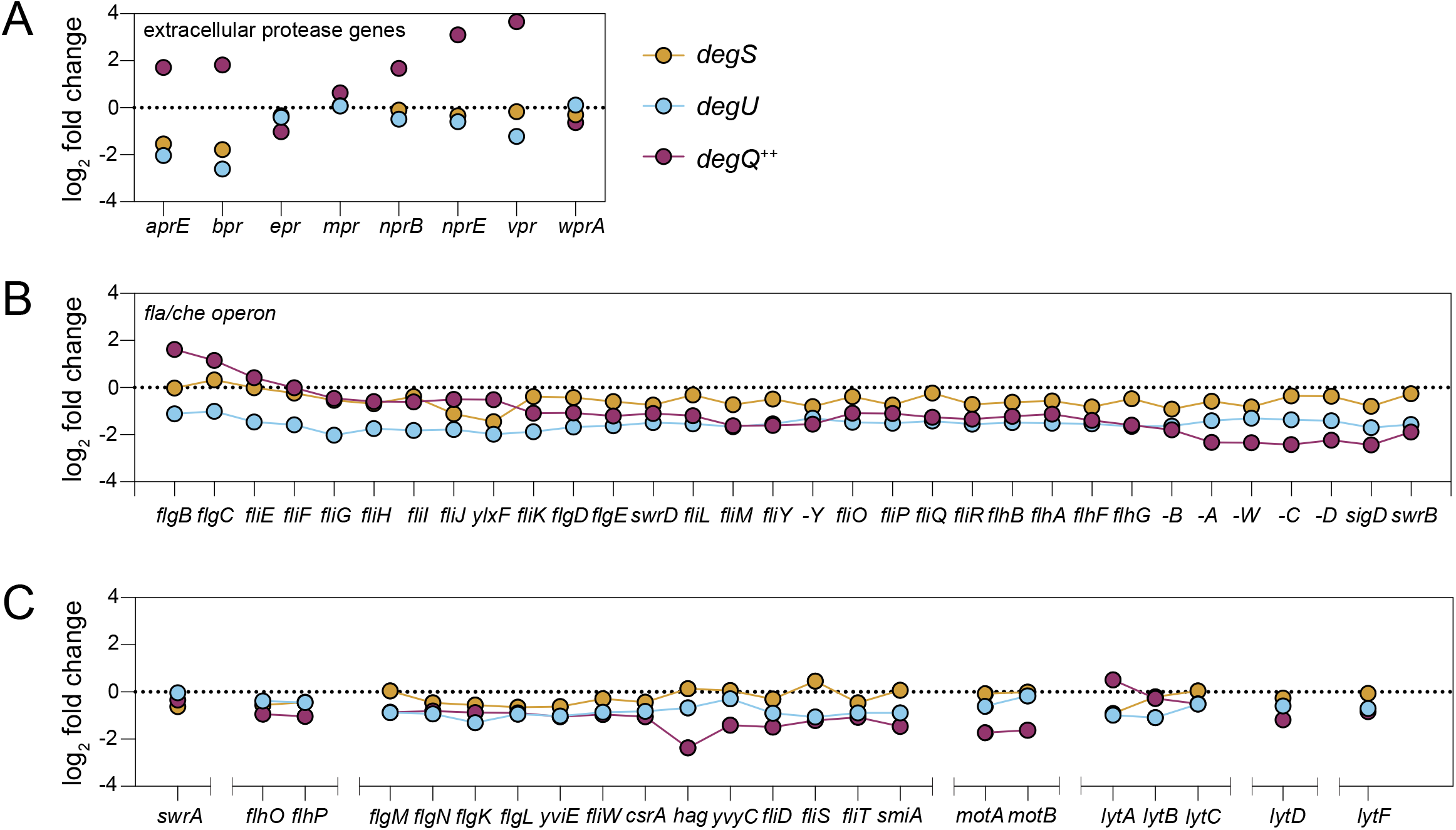
DegQ activates a subset of extracellular protease transcripts and inhibits *fla/che* operon transcript downstream of the promoter. Transcript abundance expressed as a log_2_ fold change relative to wild type for the indicated genes in the indicated strains: *degU* deletion mutant (blue, DS3649), *degS* deletion mutant (orange, DK4236), and a strain overexpressing *degQ* from a P*_hyspank_* promoter induced with 1 mM IPTG (*degQ*^++^, magenta, DK6294). Cells were collected after growth in LB to an OD_600_ of 2. Genes and operons not drawn to scale. A) Change in transcript abundance for the 8 genes known to encode extracellular proteases. B) Change in transcript abundance for each of the genes in the 32 gene *fla/che* operon. C) Change in transcript abundance for genes under SigD control.

### DegQ is an activator of extracellular proteases and an inhibitor of swarming motility

DegQ is thought to control the phosphorylation state of DegU by modulating either the kinase and/or phosphatase activity of DegS (13, 37). Consistent with a positive role on DegU-dependent phenotypes, mutation of the *degQ* gene reduced extracellular protease activity (**Fig 2**), and reduced transcript levels of both the *aprE* and *bpr* genes in transcriptomic analysis (**Fig 4A**). We note, however, that the mutation of *degQ* did not result in reduced expression of the P*_aprE_* reporter (**Fig 3A**). We conclude that while DegQ appears necessary for activating DegU-dependent expression of extracellular protease genes, it does not appear to do so at the level of promoter initiation. Rather, we infer that DegQ may act downstream of the P*_aprE_* promoter to increase the level of the *aprE* gene transcript.

To determine whether DegQ was sufficient to activate *aprE* expression, the *degQ* gene was placed under an IPTG-inducible promoter, and cells were grown in the presence of inducer (*degQ*^++^). Induction of DegQ did not cause premature expression from the P*_aprE_* reporter or extracellular protease activity, but upon entry into stationary phase, both measurements transiently increased by approximately 10-fold over wild type (**Fig 1D**, **Fig 2**). Moreover, elevated P*_aprE_* reporter expression was dramatically reduced when DegQ was induced in either a *degU* or *degS* mutant background (**Fig 3A**). Finally, RNAseq analysis of cells overexpressing DegQ indicated an increase in the transcript level of *aprE*, *bpr*, *nprB*, *nprE*, and *vpr* (**Fig 4A**). We conclude that DegQ is not essential for the expression of *aprE* but the expression of *aprE* is elevated in a DegS-DegU dependent manner when DegQ is overproduced.

To determine whether DegQ was a generalized enhancer for the DegS-DegU two component system, the expression of P*_fla/che_*, another promoter under DegU control, was measured under the same early-stationary phase conditions that was used for measuring protease gene expression. Consistent with previous reports, deletion of *degU* reduced expression from P*_fla/che_* (18, 19, 33), but here we find that deletion of either *degS* or *degQ* did not (**Fig 3B**). Moreover, artificial overexpression of DegQ did not reduce P*_fla/che_* expression in wild type, and deletion of *degU* was epistatic to *degQ* overexpression (**Fig 3B**). We conclude that DegQ is not a generalized enhancer for DegU-dependent gene expression and that transcriptional activation by DegQ is promoter-dependent. We further conclude that unlike that observed with the P*_aprE_* promoter, DegS was not required for the DegU-dependent activation of the P*_fla/che_* promoter.

The level of P*_fla/che_* promoter expression is a critical determinant of swarming motility (30, 32). Consistent with the reduced reporter activity and previous reports, deletion of *degU* (**Fig 5A**) abolished swarming (13, 14), but mutation of *degS* did not (**Fig 5B, Fig S1B**). Despite having little effect on the P*_fla/che_* reporter gene expression, deletion of *degQ* resulted in a reduced swarm lag period (**Fig 5C**), and overexpression of DegQ severely impaired swarming (**Fig 5D**). Simultaneous mutation of both *degS* and *degQ* restored a wild type swarming lag period suggesting that the *degS* mutation was epistatic (**Fig 5E**). Moreover, mutation of *degS* also abolished the inhibition of swarming due to DegQ overexpression, further supporting *degS* epistasis (**Fig 5F**). We conclude that DegQ inhibits swarming motility in a DegS-dependent manner. We infer that whatever the mechanism by which DegQ inhibits swarming, its repressive effect likely operates downstream of the P*_fla/che_*promoter.

**Figure Legend 5.**
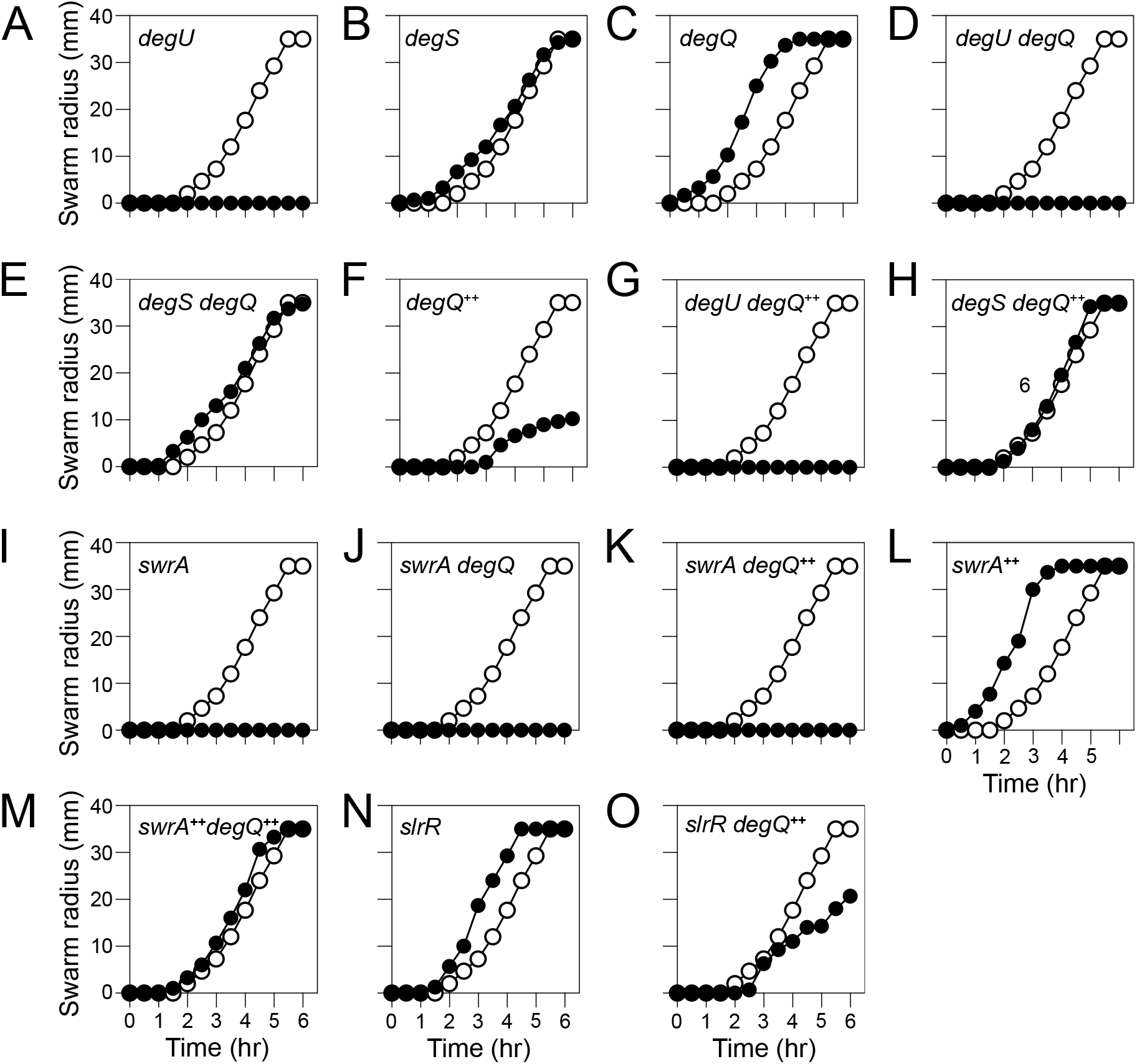
DegQ inhibits swarming motility. Swarm expansion assays for each indicated strain grown on 0.7% LB at 37 °C. In each graph, swarming of the wild type (DK1042) is represented by open symbols and swarming of the indicated mutant strain is in closed symbols. The following strains were used to generate this panel: A) *degU* (DS3649) B) *degS* (DK4236) C) *degQ* (DB3074) D) *degU degQ* (DB3744) E) *degS degQ* (DB2851) F) *degQ*^++^ (DK6294) G) *degU degQ^++^* (DB3745) H) and *degS degQ*^++^ (DB2863) I) *swrA* (DS2415) J) *swrA degQ* (DB3726) K) *swrA degQ++* (DB3185) L) *swrA^++^* (DB3835) M) *swrA^++^ degQ*^++^ (DB3729) N) *slrR* (DB3899) O) *slrR degQ^++^* (DB3910).

To determine where DegQ inhibited motility within the flagellar regulon, RNAseq transcriptomics were performed on a strain that overexpressed DegQ. Transcript levels of genes early in the *fla/che* operon were initially higher than wild type when DegQ was overexpressed, but the transcript levels gradually decreased in a distance dependent manner until the transcript levels towards the back of the operon resembled cells lacking DegU. We noted that transcript decline appeared to occur in discrete steps, reminiscent of roadblock attenuation within the *fla/che* operon recently reported to be mediated by a heteromeric complex of DNA binding proteins SinR and SlrR (40).

Disruption of SlrR in a strain is capable of swarming motility (**Fig 5G**) that overexpressed DegQ improved swarming motility but did not restore behavior to that of the wild type (**Fig 5H**). We conclude that DegQ inhibits the *fla/che* operon transcript levels downstream of initiation at the P*_fla/che_* promoter and attenuation within the operon by the SinR•SlrR heteromer contributes to the swarming deficiency.

### DegQ and SwrA are antagonistic on DegU activity

DegQ appeared to have opposing effects on various DegU-activated phenotypes. Specifically, it enhanced extracellular protease activity but inhibited swarming over surfaces. We wondered whether the differential effects were due to the presence of another DegU modulator protein, SwrA (33, 34, 41). Indeed, mutation of *swrA* resulted in a dramatic increase in the expression of P*_aprE_* and extracellular proteolytic activity and overexpression of *swrA*, reduced P*_aprE_*expression and proteolysis to levels resembling the *degU* mutant (**Fig 2A**). Moreover, the repressive effects of SwrA on both P*_aprE_* expression and extracellular protease activity were DegQ dependent, as elevated *aprE* expression was restored to wild type when both *swrA degQ* were simultaneously deleted (**Fig 2A**, **Fig 3**). Whereas SwrA appears to repress *aprE* expression indirectly through DegQ. SwrA has been shown to directly activate the P*_flache_* promoter directly, and consistent with direct activation, mutation of *swrA* was epistatic to mutation of *degQ* for P*_fla/che_* reporter activity (**Fig 2B**). We conclude that SwrA activates some members of the DegU regulon and inhibits others, and at least in the case of *aprE*, SwrA inhibition was DegQ-dependent.

One way in which SwrA might inhibit P*_aprE_* expression in a DegQ-dependent manner is if SwrA inhibits the expression of DegQ itself. To determine whether the *degQ* gene is regulated by *swrA*, a transcriptional reporter was generated in which the promoter region of *degQ* (P*_degQ_*) was fused to the *lacZ* gene and inserted at an ectopic site in the chromosome. Mutation of *swrA* resulted in a dramatic increase in the expression of *degQ* relative to wild type, consistent with SwrA functioning as a *degQ* repressor (**Fig 6**). Seemingly inconsistent with SwrA acting as a degQ repressor however, overexpression of SwrA did not decrease *degQ* expression, suggesting that the absence of SwrA and its overexpression, had two different effects depending on the promoter (**Fig 6**). We conclude that SwrA inhibits the expression of AprE indirectly, at least in part, due to the inhibition of the DegQ activator. We infer, due to the inconsistent results between the *swrA* mutant and *swrA* overexpression state, that SwrA likely inhibits *degQ* gene expression indirectly.

**Figure Legend 6.**
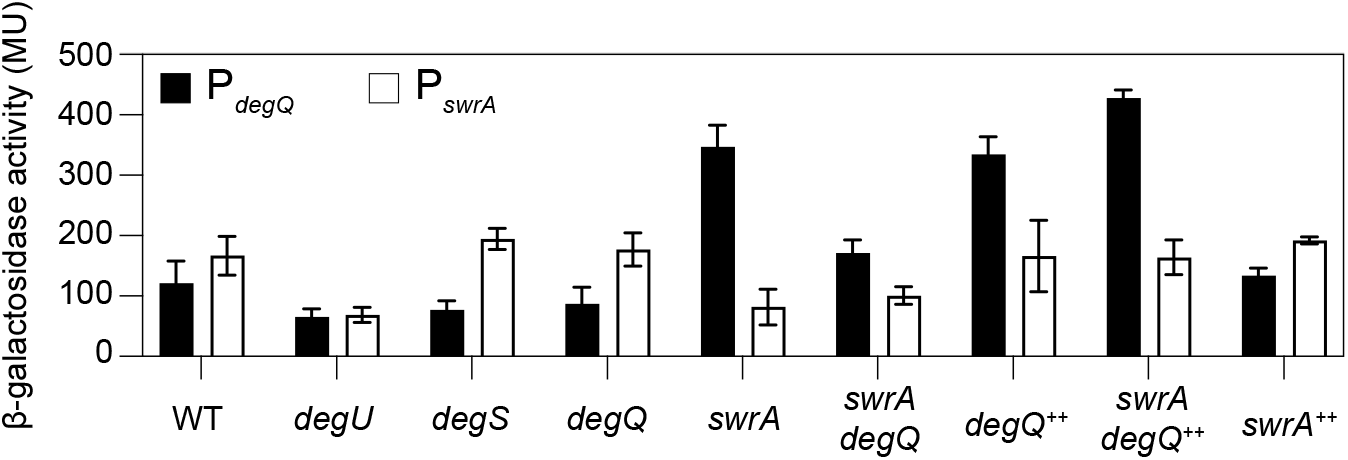
SwrA inhibits DegQ and DegQ autoactivates P*_degQ_* expression. β-galactosidase activities measured in Miller units (MU) for each of the indicated transcriptional reporter fusions and mutants grown to an OD of 2 in LB media. Expression levels from the P*_degQ_* and P*_swrA_* reporters. The following strains containing the reporter were used to generate this panel (P*_degQ_*/P*_swrA_*): wild type (DB1051/DK6613), *degU* (DB2508/DK6626), *degS* (DB3827/DB3826), *degQ* (DB2690/DB3776), *swrA* (DB3774/DB3780), *swrA degQ* (DB3959/DB3690), *degQ^++^* (DB3777/DB3778), *swrA degQ*^++^ (DB3961/ DB362), *swrA*^++^ (DB3779/DK6624). Each bar is the average of three biological replicates, and the error bars represent the standard deviation.

**Figure Legend 7.**
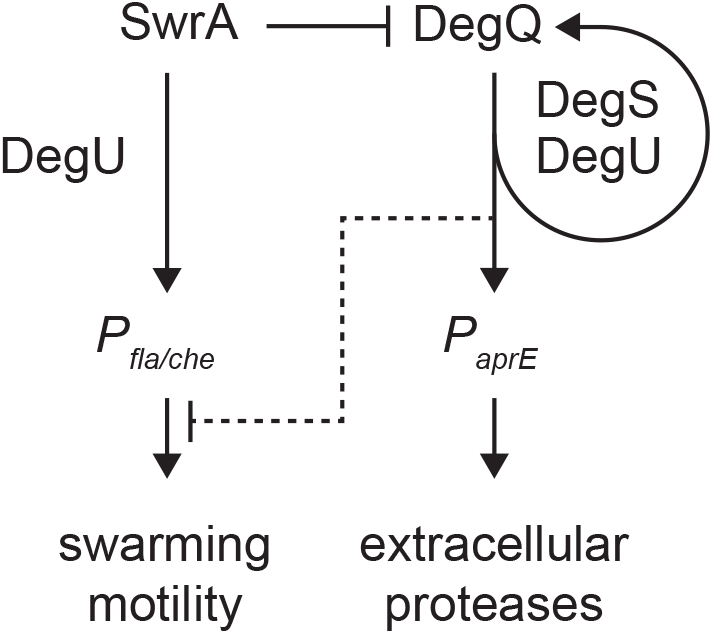
DegQ and SwrA have opposite effects on extracellular protease production and swarming motility. Schematic depicting the regulatory effects of the DegS-DegU two-component system, DegQ, and SwrA on extracellular protease production and swarming motility. Arrows indicate activation, T-bars indicate repression. Dashed T-bar indicates that DegQ inhibits swarming motility downstream of initiation at the P*_fla/che_*promoter

Transcription of the *degQ* gene has been shown to be activated by DegS and DegU (42), and mutation of either *degS* or *degU* reduced expression from the P*_degQ_*promoter. Moreover, mutation of *degU* was epistatic to mutation of *swrA* for P*_degQ_* reporter expression (**Fig 6**). We conclude that SwrA inhibits *degQ* indirectly through DegU. and was epistatic to mutation of *swrA*, indicating that inhibition by SwrA was DegU-dependent. As DegQ is also a modulator of DegU-dependent gene expression we wondered whether DegQ was autoregulatory. Indeed, mutation and overexpression of *degQ*, decreased and increased *degQ* gene expression, respectively. While SwrA caused DegU-dependent repression of the P*_degQ_* promoter, the reverse was not true, that is, mutation and overexpression seemed to have little effect on the DegU-dependent expression of the P*_swrA_* promoter. We conclude that two different modulators of DegU differential effects on the DegU regulon, and that SwrA antagonizes the production of DegQ.

## DISCUSSION

DegU is a highly-pleiotropic and poorly-understood regulator in *Bacillus subtilis* expressed with, and phosphorylated by, the cognate soluble histidine kinase DegS, but the input to DegS activity and the output DegU DNA binding site is largely unknown.

Perhaps one reason for the difficulty in resolving the function of DegU is the extensive use of domesticated laboratory strains where null mutants have little phenotypic effect and data is complicated by the reliance on hyperactive DegU mutant alleles (e.g. DegUhy32^H12L^) (9, 24, 43). Phenotypes have been reported for *degU* null mutants in ancestral strains however, likely because ancestral strains are functional for two small proteins, DegQ and SwrA, that enhance DegU activity (13, 14, 41, 43). Here we study the relative roles of DegQ and SwrA in the ancestral strain background of *B. subtilis* and find that the two proteins act in opposition to inversely activate extracellular protease production and flagellar motility, respectively.

DegQ is a 46 amino acid protein, the expression of which was abolished in domesticated strains by mutations in the promoter (35, 44, 45). Consistent with early genetics, we find that DegQ increases both extracellular protease activity and the expression of *aprE* (35, 46). How DegQ activates gene expression is unknown, save that it requires both DegS and DegU, and enhances DegS-dependent DegU phosphorylation *in vitro* (13, 42, 46). AlphaFold3 modeling suggests that DegQ may interact with the N-terminal input domain of DegS (37), and DegQ likely either potentiates DegS signaling by altering its conformation, or perhaps DegQ is the signal itself. Whatever the case, overexpression of DegQ is insufficient to activate *aprE* gene expression until the transition to stationary phase and this may explain the absence of DegU binding to the *aprE* promoter region in ChIP-seq analysis of cells during exponential growth (33). We infer that extracellular protease expression is somehow restricted, and *aprE* is known to be inhibited until the onset of stationary phase by the multiple repressors including ScoC and CodY (47–50).

DegQ also functions as an inhibitor of swarming motility as here we show that mutation of *degQ* decreases the swarming lag period while the overexpression of *degQ* impairs swarm expansion. Swarming has been shown to depend on a critical density of flagella on the cell surface (32) and flagellar number has been shown to be proportional to the level of expression from the P*_flache_*promoter (30). DegQ does not seem to inhibit P*_fla/che_* promoter activity however and instead causes a gradual decrease in transcript abundance from the front of the *fla/che* operon to the back. The *fla/che* operon was recently reported to experience transcript attenuation due to internal binding of the SinR•SlrR a heteromer that seems to cause premature termination (40), and mutation of SlrR improved motility when DegQ was overexpressed. Thus, it seems that DegQ inhibits motility, at least in part, by enhancing the amount and/or activity of the SinR•SlrR heteromeric repressor and we suspect that the DegUhy32 allele likely inhibits motility by the same mechanism (17, 40). How DegQ activates heteromer activity is unknown.

SwrA is a 117 amino acid protein, the expression of which was abolished in domesticated strains by an early loss-of-function frameshift mutation in the coding region (27, 28, 44). SwrA increases the frequency of motile cells in the population, increases the number of flagella per cell, and promotes swarming motility (27, 30). Previous work has shown that SwrA interacts with the DegU N-terminal domain and causes DegU oligomerization to activate expression of the P*_fla/che_* promoter (33, 34). Here we find that SwrA inhibits expression of the P*_aprE_* promoter indirectly by repressing expression of the *degQ* gene. How SwrA acts as a repressor is unknown but, in its absence, DegQ autoactivates its DegU-dependent expression, likely driving an increase in DegQ abundance. As the frameshift mutation in *swrA* is one of the earliest events in *B. subtilis* domestication, we wonder whether the promoter down-mutation in *degQ* was selected as a consequence of the absence of SwrA that would otherwise prevent DegQ positive feedback (44, 51). Thus, SwrA promotes flagellar gene expression during exponential growth, and we infer that during the transition to stationary phase, SwrA is inactivated thereby derepressing DegQ and extracellular protease gene expression.

In sum, we find that two small proteins SwrA and DegQ fundamentally alter the activity of the DegU response regulator. SwrA promotes activation of the *P_fla/che_*promoter by mechanism that depends on DegU but not DegS. By contrast, DegQ activates the expression of *aprE* in a manner that depends on both components of the DegS/DegU two-component system. Moreover, while SwrA directly inhibits DegQ production, DegQ overrides SwrA function downstream of its target site and attenuates the production of new flagella during stationary phase. From the combined action of the two proteins, we infer that de novo flagellar synthesis and the secretion of extracellular proteases do not occur simultaneously, at least at the level of an individual cell. It is unclear why *B. subtilis* would evolve multiple modulators of a single two-component system rather than evolve an alternative regulator, but the fact that it did may explain why the *degU* regulon seems both vast and mysterious. We wonder whether the various modulators not only govern the timing of gene expression within the regulon, but might also fundamentally alter the DegU binding site, thereby obscuring consensus.

## MATERIALS AND METHODS

### Growth conditions

*B. subtilis* stains were grown in Luria-Bertani (LB) broth (10 g tryptone, 5 g yeast extract, 5 g NaCl per L), or LB plates fortified with 1.5% Bacto agar at 37 °C. When appropriate, antibiotics were included in the following concentrations: 5 µg/mL chloramphenicol (cat), 5 µg/mL kanamycin (kan), 1 µg/mL erythromycin plus 25 µg/mL lincomycin (mls), 100 µg/mL spectinomycin (spec), 10 µg/mL tetracycline (tet). Isopropyl β-D-thiogalactopyranoside (IPTG, Sigma) was added to the medium at 1 mM when appropriate.

### SPP1 phage transduction

Serial dilutions of SPP1 phage were added to 200 µL of dense culture grown in TY broth (LB broth supplemented after autoclaving with 10 mM MgSO_4_ and 100 µM MnSO_4_). To each mixture, 3 mL of molten TY top agar (TY broth supplemented with 0.5% agar) was added. The mixture poured atop fresh TY plates (TY broth fortified with 1.5% agar), and the plates were incubated at 30 °C overnight. Top agar from a plate containing near-confluent plaques was harvested by scraping into a 15 mL conical tube with 5 mL of TY broth and centrifuged at 6,000 x g for 5 min. The supernatant was passed through a 0.22 µm syringe filter and stored at 4 °C. Recipient cells for transduction were grown to stationary phase in 3 mL TY broth at 37 °C. One milliliter of dense culture was mixed with 25 µL of SPP1 donor phage stock made above. Nine milliliters of TY broth were added and mixed by rocking at room temperature for 30 min. The transduction mixture was centrifuged at 6,000 x g for 5 min, the supernatant was discarded, and the pellet was suspended in the remaining volume. 100 µL of cell suspension was then plated on LB plates fortified with the appropriate antibiotic for selection and 10 mM sodium citrate if the antibiotic was cat, kan, or spec.

### Strain construction

All constructs were either first introduced into the domesticated *B. subtilis* strain PY79 by natural competence and then transferred to the undomesticated *B. subtilis* 3610 background using SPP1-mediated generalized phage transduction (52) or transformed directly into DK1042 (53). All plasmids used in this study are listed in Table S1. All primers used in this study are listed in Table S2. All strains used in this study are listed in Table 1.

**Table 1:**
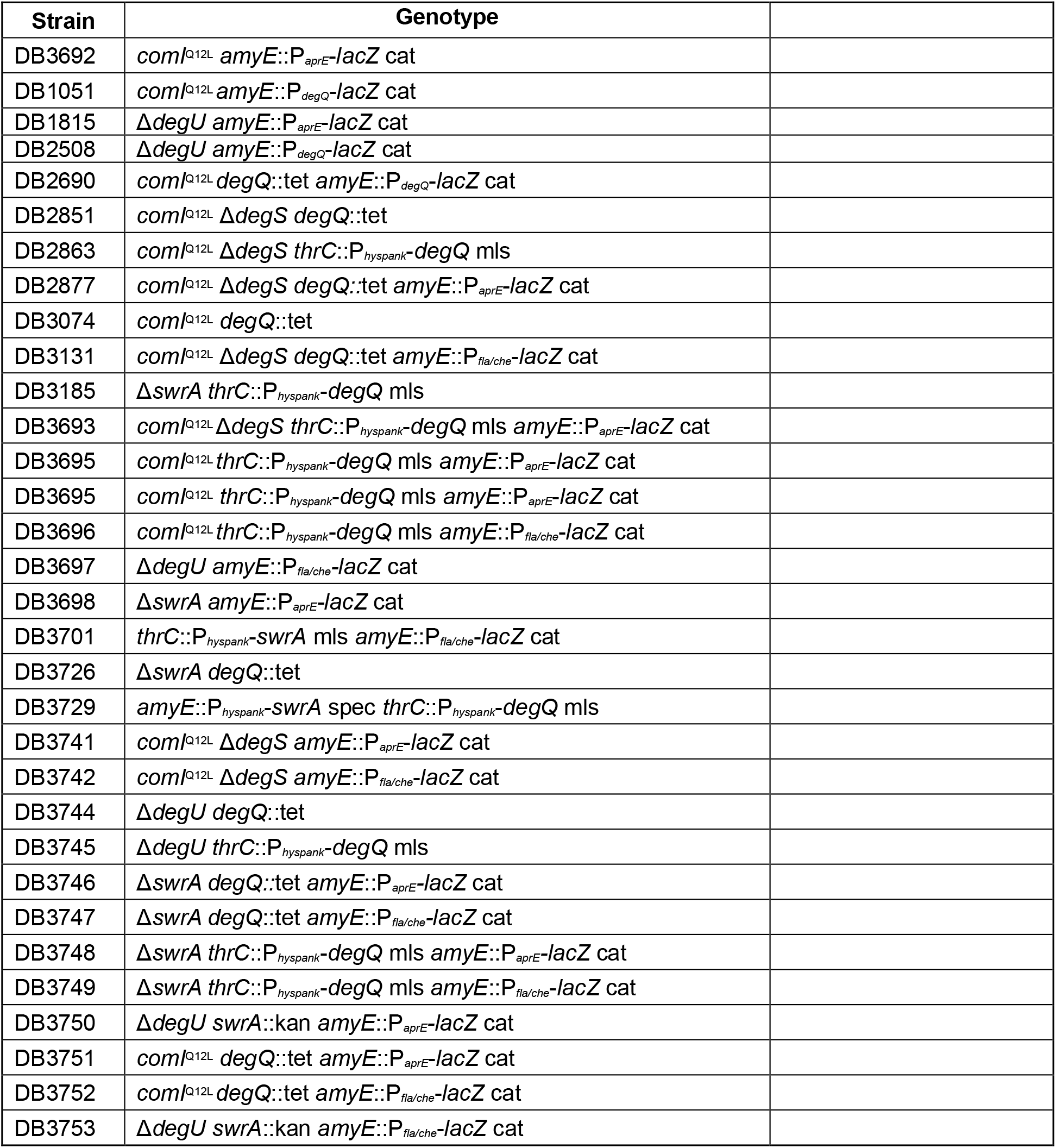
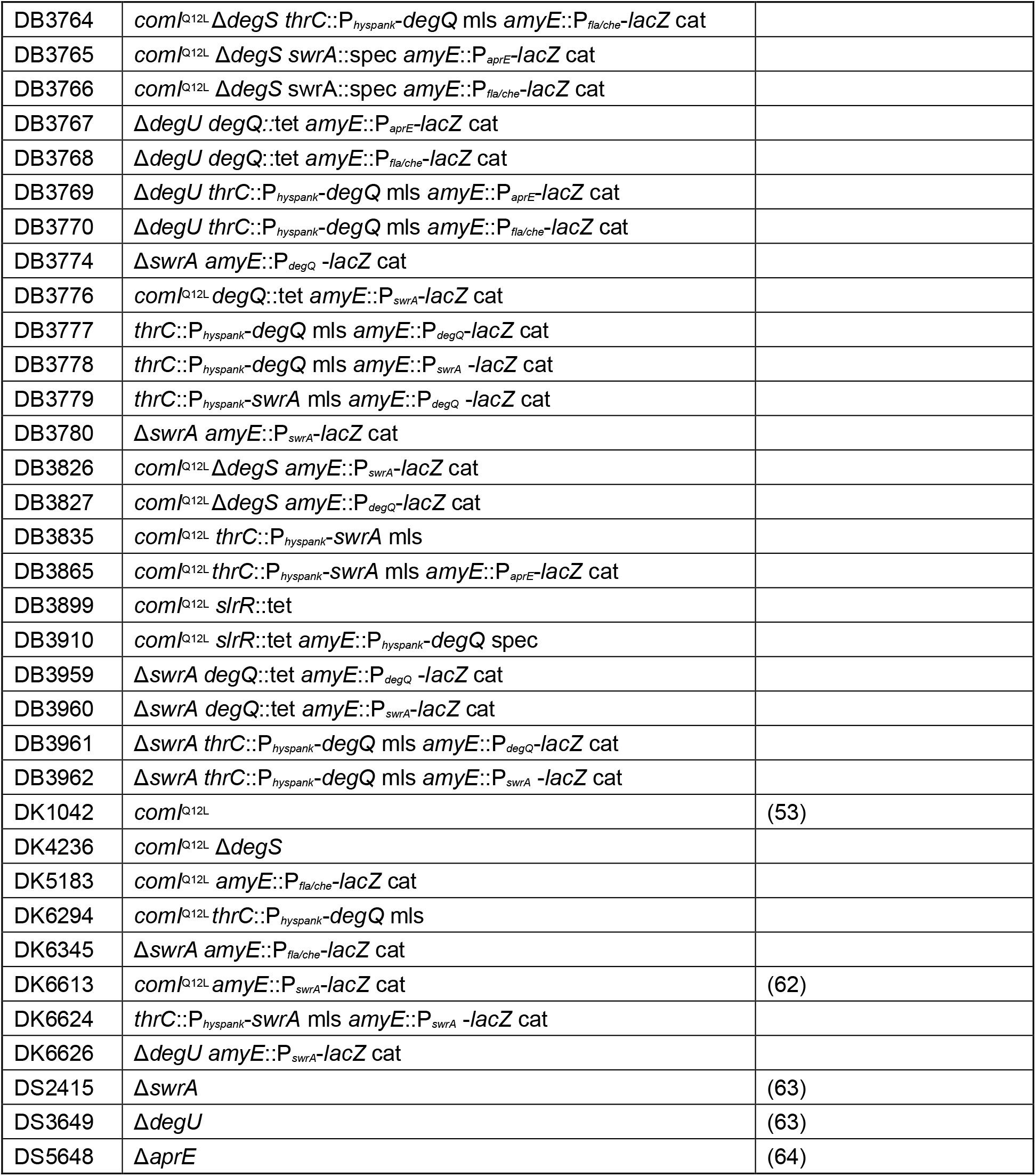
Strains.

| Strain | Genotype |  |
| --- | --- | --- |
| DB3692 | *comI*<sup>Q12L</sup> *amyE*::P<sub>aprE</sub>-*lacZ* cat |  |
| DB1051 | *comI*<sup>Q12L</sup> *amyE*::P<sub>degQ</sub>-*lacZ* cat |  |
| DB1815 | Δ*degU* *amyE*::P<sub>aprE</sub>-*lacZ* cat | |
| DB2508 | Δ*degU* *amyE*::P<sub>degQ</sub>-*lacZ* cat | |
| DB2690 | *comI*<sup>Q12L</sup> *degQ*::tet *amyE*::P<sub>degQ</sub>-*lacZ* cat |  |
| DB2851 | *comI*<sup>Q12L</sup> Δ*degS* *degQ*::tet | |
| DB2863 | *comI*<sup>Q12L</sup> Δ*degS* *thrC*::P<sub>hyspank</sub>-*degQ* mls | |
| DB2877 | *comI*<sup>Q12L</sup> Δ*degS* *degQ*::tet *amyE*::P<sub>aprE</sub>-*lacZ* cat | |
| DB3074 | *comI*<sup>Q12L</sup> *degQ*::tet |  |
| DB3131 | *comI*<sup>Q12L</sup> Δ*degS* *degQ*::tet *amyE*::P<sub>fla/che</sub>-*lacZ* cat | |
| DB3185 | Δ*swrA* *thrC*::P<sub>hyspank</sub>-*degQ* mls | |
| DB3693 | *comI*<sup>Q12L</sup> Δ*degS* *thrC*::P<sub>hyspank</sub>-*degQ* mls *amyE*::P<sub>aprE</sub>-*lacZ* cat | |
| DB3695 | *comI*<sup>Q12L</sup> *thrC*::P<sub>hyspank</sub>-*degQ* mls *amyE*::P<sub>aprE</sub>-*lacZ* cat |  |
| DB3695 | *comI*<sup>Q12L</sup> *thrC*::P<sub>hyspank</sub>-*degQ* mls *amyE*::P<sub>aprE</sub>-*lacZ* cat |  |
| DB3696 | *comI*<sup>Q12L</sup> *thrC*::P<sub>hyspank</sub>-*degQ* mls *amyE*::P<sub>fla/che</sub>-*lacZ* cat |  |
| DB3697 | Δ*degU* *amyE*::P<sub>fla/che</sub>-*lacZ* cat | |
| DB3698 | Δ*swrA* *amyE*::P<sub>aprE</sub>-*lacZ* cat | |
| DB3701 | *thrC*::P<sub>hyspank</sub>-*swrA* mls *amyE*::P<sub>fla/che</sub>-*lacZ* cat |  |
| DB3726 | Δ*swrA* *degQ*::tet | |
| DB3729 | *amyE*::P<sub>hyspank</sub>-*swrA* spec *thrC*::P<sub>hyspank</sub>-*degQ* mls |  |
| DB3741 | *comI*<sup>Q12L</sup> Δ*degS* *amyE*::P<sub>aprE</sub>-*lacZ* cat | |
| DB3742 | *comI*<sup>Q12L</sup> Δ*degS* *amyE*::P<sub>fla/che</sub>-*lacZ* cat | |
| DB3744 | Δ*degU* *degQ*::tet | |
| DB3745 | Δ*degU* *thrC*::P<sub>hyspank</sub>-*degQ* mls | |
| DB3746 | Δ*swrA* *degQ*::tet *amyE*::P<sub>aprE</sub>-*lacZ* cat | |
| DB3747 | Δ*swrA* *degQ*::tet *amyE*::P<sub>fla/che</sub>-*lacZ* cat | |
| DB3748 | Δ*swrA* *thrC*::P<sub>hyspank</sub>-*degQ* mls *amyE*::P<sub>aprE</sub>-*lacZ* cat | |
| DB3749 | Δ*swrA* *thrC*::P<sub>hyspank</sub>-*degQ* mls *amyE*::P<sub>fla/che</sub>-*lacZ* cat | |
| DB3750 | Δ*degU* *swrA*::kan *amyE*::P<sub>aprE</sub>-*lacZ* cat | |
| DB3751 | *comI*<sup>Q12L</sup> *degQ*::tet *amyE*::P<sub>aprE</sub>-*lacZ* cat |  |
| DB3752 | *comI*<sup>Q12L</sup> *degQ*::tet *amyE*::P<sub>fla/che</sub>-*lacZ* cat |  |
| DB3753 | Δ*degU* *swrA*::kan *amyE*::P<sub>fla/che</sub>-*lacZ* cat | |
| DB3764 | *comI*<sup>Q12L</sup> Δ*degS* *thrC*::P<sub>hyspank</sub>-*degQ* mls *amyE*::P<sub>fla/che</sub>-*lacZ* cat | |
| DB3765 | *comI*<sup>Q12L</sup> Δ*degS* *swrA*::spec *amyE*::P<sub>aprE</sub>-*lacZ* cat | |
| DB3766 | *comI*<sup>Q12L</sup> Δ*degS* *swrA*::spec *amyE*::P<sub>fla/che</sub>-*lacZ* cat | |
| DB3767 | Δ*degU* *degQ*::tet *amyE*::P<sub>aprE</sub>-*lacZ* cat | |
| DB3768 | Δ*degU* *degQ*::tet *amyE*::P<sub>fla/che</sub>-*lacZ* cat | |
| DB3769 | Δ*degU* *thrC*::P<sub>hyspank</sub>-*degQ* mls *amyE*::P<sub>aprE</sub>-*lacZ* cat | |
| DB3770 | Δ*degU* *thrC*::P<sub>hyspank</sub>-*degQ* mls *amyE*::P<sub>fla/che</sub>-*lacZ* cat | |
| DB3774 | Δ*swrA* *amyE*::P<sub>degQ</sub> -*lacZ* cat | |
| DB3776 | *comI*<sup>Q12L</sup> *degQ*::tet *amyE*::P<sub>swrA</sub>-*lacZ* cat |  |
| DB3777 | *thrC*::P<sub>hyspank</sub>-*degQ* mls *amyE*::P<sub>degQ</sub>-*lacZ* cat |  |
| DB3778 | *thrC*::P<sub>hyspank</sub>-*degQ* mls *amyE*::P<sub>swrA</sub> -*lacZ* cat |  |
| DB3779 | *thrC*::P<sub>hyspank</sub>-*swrA* mls *amyE*::P<sub>degQ</sub> -*lacZ* cat |  |
| DB3780 | Δ*swrA* *amyE*::P<sub>swrA</sub>-*lacZ* cat | |
| DB3826 | *comI*<sup>Q12L</sup> Δ*degS* *amyE*::P<sub>swrA</sub>-*lacZ* cat | |
| DB3827 | *comI*<sup>Q12L</sup> Δ*degS* *amyE*::P<sub>degQ</sub>-*lacZ* cat | |
| DB3835 | *comI*<sup>Q12L</sup> *thrC*::P<sub>hyspank</sub>-*swrA* mls |  |
| DB3865 | *comI*<sup>Q12L</sup> *thrC*::P<sub>hyspank</sub>-*swrA* mls *amyE*::P<sub>aprE</sub>-*lacZ* cat |  |
| DB3899 | *comI*<sup>Q12L</sup> *slrR*::tet |  |
| DB3910 | *comI*<sup>Q12L</sup> *slrR*::tet *amyE*::P<sub>hyspank</sub>-*degQ* spec |  |
| DB3959 | Δ*swrA* *degQ*::tet *amyE*::P<sub>degQ</sub> -*lacZ* cat | |
| DB3960 | Δ*swrA* *degQ*::tet *amyE*::P<sub>swrA</sub>-*lacZ* cat | |
| DB3961 | Δ*swrA* *thrC*::P<sub>hyspank</sub>-*degQ* mls *amyE*::P<sub>degQ</sub>-*lacZ* cat | |
| DB3962 | Δ*swrA* *thrC*::P<sub>hyspank</sub>-*degQ* mls *amyE*::P<sub>swrA</sub> -*lacZ* cat | |
| DK1042 | *comI*<sup>Q12L</sup> | (53) |
| DK4236 | *comI*<sup>Q12L</sup> Δ*degS* | |
| DK5183 | *comI*<sup>Q12L</sup> *amyE*::P<sub>fla/che</sub>-*lacZ* cat |  |
| DK6294 | *comI*<sup>Q12L</sup> *thrC*::P<sub>hyspank</sub>-*degQ* mls |  |
| DK6345 | Δ*swrA* *amyE*::P<sub>fla/che</sub>-*lacZ* cat | |
| DK6613 | *comI*<sup>Q12L</sup> *amyE*::P<sub>swrA</sub>-*lacZ* cat | (62) |
| DK6624 | *thrC*::P<sub>hyspank</sub>-*swrA* mls *amyE*::P<sub>swrA</sub> -*lacZ* cat |  |
| DK6626 | Δ*degU* *amyE*::P<sub>swrA</sub>-*lacZ* cat | |
| DS2415 | Δ*swrA* | (63) |
| DS3649 | Δ*degU* | (63) |
| DS5648 | Δ*aprE* | (64) |

#### slrR::tet

The *slrR::tet* allele was a generous gift from Frances Chu (Harvard University).

#### degQ::tet

To generate the *degQ::tet* insertion deletion construct, DK1042 chromosomal DNA was used as a template to PCR amplify the region upstream of the *degQ* gene with primer pair 8506/8507 and the region downstream of the *degQ* gene with primer pair 8508/8509. Next, the tetracycline resistance cassette was amplified from plasmid pDG1515 (54) using primer pair 3250/3251. The three fragments were ligated together using isothermal assembly (55) and amplified with primer pair 8506/8509 prior to transformation into DK1042.

#### ΔdegS

To generate the *ΔdegS* in frame marker-less deletion construct, the region upstream of *degS* was PCR amplified using the primer pair 5172/5173, the region downstream of *degS* was PCR amplified using the primer pair 5170/5171, and the two fragments were combined and ligated by isothermal assembly. Next, the isothermal assembly product was used as a PCR template and amplified using primers 5172/5171, the product was digested with BamHI and KpnI and ligated into the BamHI and KpnI sites of the plasmid pMiniMAD (56) which carries a temperature sensitive origin of replication and an erythromycin resistance cassette to generate pAC001. The plasmid pAC001 was purified from the *recA+ E. coli* strain TG1, transformed into DK1042, plated on LB plates containing mls antibiotic, and incubated at the permissive temperature of 30 °C. An mls resistant colony was struck on a LB mls plate and incubated overnight at the non-permissive temperature of 37 °C to force single crossover integration into the chromosome. Next a colony was incubated in 3 mL LB broth overnight at room temperature and a serially dilution was plated on LB agar at 37 °C. Individual colonies were patched on LB plates and LB plates containing *mls* to identify *mls* sensitive colonies that had evicted the plasmid. Chromosomal DNA from colonies that had excised the plasmid was purified and screened by PCR using primers 5172/5171 to determine which isolate had retained the *ΔdegS* allele.

### Complementation constructs

To generate the *degS* complementation construct pDP486, the *degS* gene was amplified from DK1042 chromosomal DNA using primer pair 5299/5301, the amplicon was digested with BamHI/SalI and cloned into the BamHI/SalI sites of plasmid pAH25 containing a spectinomycin resistance cassette and polylinker between two arms of the *amyE* gene (generous gift of Amy Camp, Mount Holyoke College). To generate the *degU* complementation construct pDP487, the *degU* gene was amplified from DK4326 *ΔdegS* chromosomal DNA using primer pair 5299/5301, the amplicon was digested with BamHI/SalI and cloned into the BamHI/SalI sites of plasmid pAH25.

### Inducible constructs

To generate the inducible *degQ* construct pEC20, the *degQ* gene was amplified from DK1042 chromosomal DNA with primer pair 2957/2958, the amplicon was digested with HindIII/NheI and cloned into the HinDIII/NheI sites of pDR111 containing a spectinomycin resistance cassette, the *lacI* gene, and a polylinker downstream of the *P_hyspank_* promoter between the arms of the *amyE* gene (generous gift of David Rudner, Harvard Medical School).

To generate the inducible *degQ* construct pAC016, the *degQ* gene was amplified from DK1042 chromosomal DNA with primer pair 5310/5311, the amplicon was digested with NheI/SphI, and cloned into the NheI/SphI sites of pDP150 containing an *mls* resistance cassette, the *lacI* gene, and a polylinker downstream of the P*_hyspank_* promoter between the arms of the *thrC* gene (57).

### Transcriptional reporter constructs

To generate the β*-*galactosidase (*lacZ*) reporter constructs pDP369 and pAEJ15, PCR products containing the following promoters were amplified for *B. subtilis* DK1042 chromosomal DNA using the primers indicated in parentheses: P*_aprE_* (8442/8443), and P*_degQ_* (8119/8120). Each PCR product was digested with EcoRI and either BamHI or HindIII and cloned independently into the EcoRI and either BamHI or HindIII sites of plasmid pDG268, which carries a chloramphenicol resistance marker and a polylinker upstream of the *lacZ* gene between two arms of the *amyE* gene (58).

### β-galactosidase assays

Three biological replicates of each strain were grown in LB broth at 37 °C to an OD_600_ of 1.7-2.2. Two hundred microliters of culture were pelleted and resuspended in 200 µL of Z-buffer (16.1 g Na_2_HPO_4_•7H_2_O + 5.5 g NaH_2_PO_4_• H_2_O + 0.75 g KCl + 0.246 g MgSO_4_•7H_2_O in 1 L H2O, pH = 7) with 0.27% β-mercaptoenthanol. Four microliters of 10 mg/mL lysozyme (Dot Scientific) were added, and samples were incubated at 30 °C for 15 min to lyse. In a 96-well plate, lysates were diluted in Z-buffer to 200 µL depending on the strength of the promoter fused to lacZ (P*_aprE_* = 1:1 [undiluted], P*_fla/che_* = 1:10, P*_degQ_* = 1:1, P*_swrA_* = 1:2). Three negative control wells contained only 200 µL of Z-buffer. Forty microliters of 4 mg/mL ortho-nitrophenyl-β-galactopyranoside in Z-buffer was added to each well. The plate was incubated at 30 °C shaking in a double orbital pattern, and the OD420 and OD550 were collected every 2 min. The slope of each sample’s ODs over time was derived. The average slope of the three negative control wells was subtracted from the slope of each experimental trial to control for background. The Miller Units were calculated using the following formula:

### Protease assays

In biological triplicates, cells were inoculated in LB, supplemented with IPTG when appropriate, and incubated rolling at 37 °C until OD_600_ = 1.7-2.2. One milliliter of culture was centrifuged at 6000 x g for 10 min, and 900 µL of supernatant was collected in a fresh tube and incubated with 400 µL of 1% azocasein pH = 8.0 for 30 min at 37 °C. Each reaction was quenched by the addition of 600 µL of 10% trichloroacetic acid and incubated on ice for 30 min. Cells were centrifuged at 16,000 rpm for 5 min, and 800 µL of supernatant was added to 200 µL of 1.8 N NaOH in a clean cuvette. Absorbance was measured at OD_440_, and protease activity was calculated by dividing OD_440_/OD_600_.

### RNA extraction and sequencing

RNA was extracted from *B. subtilis* cultures grown in LB to an OD600 of ∼2.0. Five mL of culture was flash frozen by pre-chilled methanol in an equal volume and centrifuged at 5000 *g* for 10 min at 4 °C. Supernatant was removed and the pellet was resuspended in 800 µL of LETS buffer (10 mM Tris-HCl pH 7.4, 50 mM LiCl, 20 mM EDTA pH 8.0, 1% SDS) heated to 75 °C. The mixture was added to 650 mg acid-washed glass beads and 600 µL of heated acid-saturated phenol pH 4, and vortexed for 3 min. 600 µL of chloroform was added and the mixture was vortexed for 30 sec before being centrifuged at 3200 g for 10 min at 4 °C. Equal parts of aqueous phase and isopropanol were combined, inverted, and incubated at room temperature for 10 min before centrifugation at 6000 x g for 25 min at 4 °C. Supernatant was discarded, and the pellet was resuspended in a mL of cold 75% ethanol and centrifuged for 5 min at 4 °C. The resuspension was centrifuged at max 6000 x g for 5 min at 4 °C, supernatant removed, pellet air-dried and rehydrated in 20 µL nuclease-free water. Samples were heated for 5 min at 55 °C to encourage resuspension before 300 µL of TRIzol was added, and vortexed for 15 sec. Samples incubated at room temperature for 5 min and combined with 60 µL chloroform. The mixture was inverted and incubated at room temperature for 2 min before centrifugation at max speed for 15 min at 4 °C. The steps starting with the collection of the aqueous layer through the 5 min incubation at 55 °C were repeated. The RNA was treated with RNase-free DNase I at 37 °C for 30 min (Invitrogen) according to the manufacturer’s instructions and RNA was re-extracted via the process above. rRNA depletion, library preparation and analysis were performed by SeqCoast Genomics (Portsmouth, New Hampshire, USA). Illumina paired-end sequencing of the libraries was performed, and at least 12 million paired-end reads were obtained for each sample. Transcript per kilobase million were quantified using Salmon (59) and differential gene expression was calculated with DESeq2 (60) using the NCBI 3610 genome (61). Differentially expressed genes were considered significant with a P-value < 0.05 and a [log_2_(fold-change)] > 2.0.

### Quantitative swarm expansion assays

Cells were grown overnight at room temperature in LB broth, back diluted and grown to mid-log phase at 37 °C in LB broth supplemented with 1 mM IPTG when applicable and resuspended to an OD600 of 10 in MQ H_2_O containing 0.5% India ink. Freshly prepared LB plates containing 0.7% Bacto agar (25 mL/plate) and IPTG (if applicable) were dried by laminar flow hood for 10 min, centrally inoculated with 10 µL of the cell suspension in ink, dried for another 10 min, and incubated at 37 °C for 6 h. Each strain was surveyed in technical triplicate, and measurements of swarm expansion in millimeters were collected every 30 min. The India ink demarcates the origin of inoculation and for consistency an axis was drawn on the back of the plate and measurements were taken along this axis.

## ACKNOWLEDGEMENTS

We are grateful to Alex Canzoneri and Abby Jackson for plasmid construction. The work was funded by NIH grant R35 GM131783 to D.B.K.

---

**Table S1:**

| Plasmid | Genotype |
| --- | --- |
| pDP639 | *amyE*::P<sub>aprE</sub>-*lacZ* cat *amp* |
| pAH25 | *amyE*::spec *amp* |
| pAC001 | $\Omega\Delta degS$ *ori<sup>TsBs</sup> mls amp* |
| pAC016 | *thrC::P<sub>hyspank</sub>-degQ mls amp* |
| pAEJ15 | *amyE::P<sub>degQ</sub>-lacZ cat amp* |
| pDG1515 | *tet amp* |
| pDG268 | *amyE::lacZ cat amp* |
| pDP150 | *thrC::P<sub>hyspank</sub> mls amp* |
| pDP486 | *amyE::P<sub>degS</sub>-degS spec amp* |
| pDP487 | *amyE::P<sub>degS</sub>-degU spec amp* |
| pDR111 | *amyE::P<sub>hyspank</sub> spec amp* |
| pEC20 | *amyE::P<sub>hyspank</sub>-degQ spec amp* |
| pMiniMAD | *ori<sup>TsBs</sup> mls amp* |

**Table S2:**
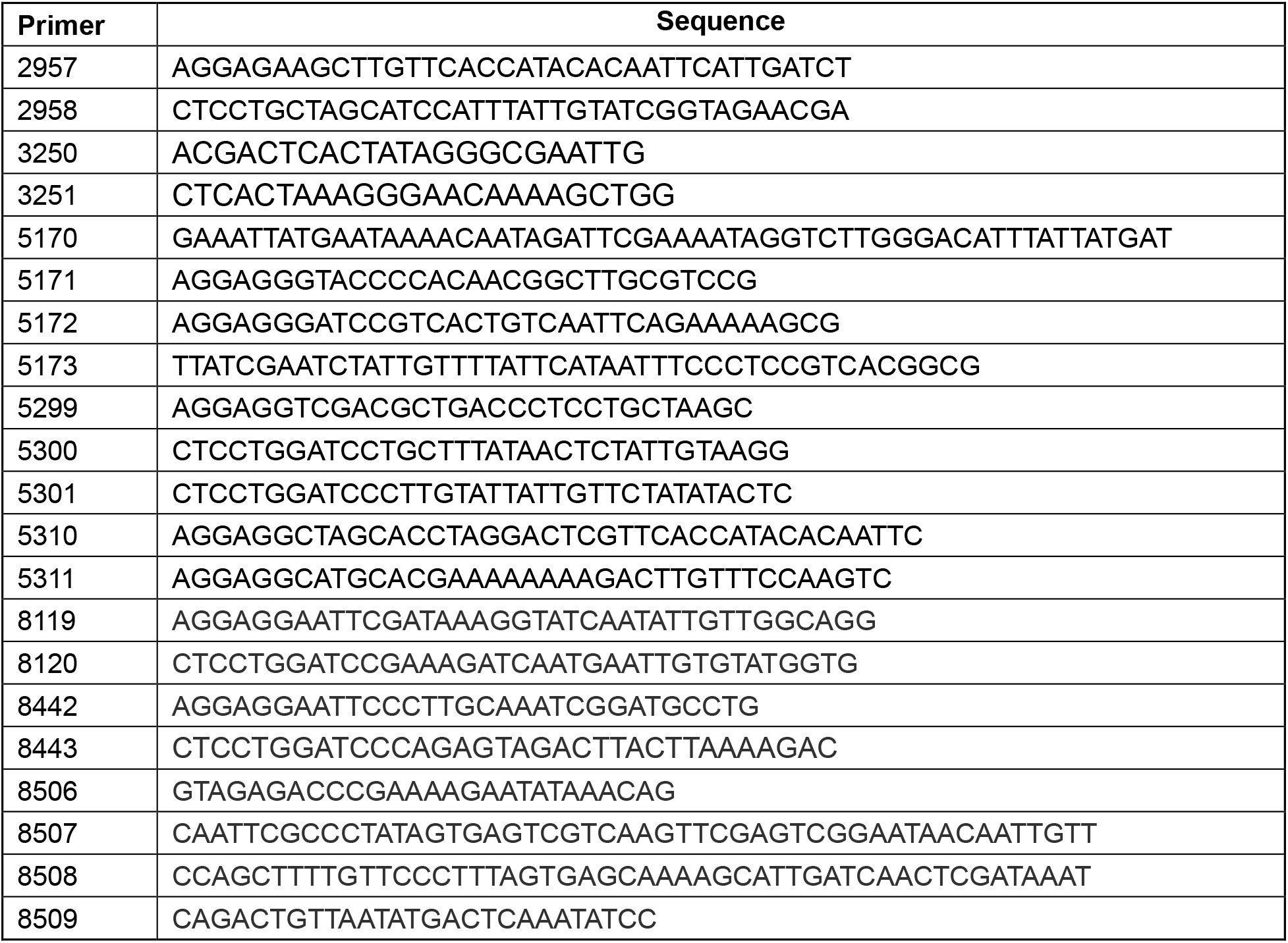

